# Presynaptic Endoplasmic Reticulum Contributes Crucially to Short-term Plasticity in Small Hippocampal Synapses

**DOI:** 10.1101/431866

**Authors:** Nishant Singh, Thomas Bartol, Herbert Levine, Terrence Sejnowski, Suhita Nadkarni

**Affiliations:** Division of Biology, Indian Institue of Science Education and Research, Dr. Homi Bhabha Road, Pune, India; Computational Neurobiology Laboratory, Salk Institute for Biological Studies, 10010 N Torrey Pines Rd, La Jolla, CA 92037; Center for Theoretical and Biological Physics, Rice University, Houston, TX 77005-1827, United States of America

## Abstract

Short-term plasticity (STP) of the presynaptic terminal maintains a brief history of activity experienced by the synapse that may otherwise remain unseen by the postsynaptic neuron. These synaptic changes are primarily regulated by calcium dynamics in the presynaptic terminal. A rapid increase in intracellular calcium is initiated by the opening of voltage-dependent calcium channels in response to depolarization, the main source of calcium required for vesicle fusion. Separately, electron-microscopic studies of hippocampal CA3-CA1 synapses reveal the strong presence of endoplasmic reticulum (ER) in all presynaptic terminals. However, the precise role of the ER in modifying STP at the presynaptic terminal remains unexplored. To investigate the contribution of ER in modulating calcium dynamics in small hippocampal boutons, we performed *in silico* experiments in a physiologically-realistic canonical synaptic geometry based on reconstructions of CA3-CA1 Schaffer collaterals in the rat hippocampus. The model predicts that presynaptic calcium stores are critical in generating the observed paired-pulse ratio (PPR) of normal CA3-CA1 synapses. In control synapses with intact ER, SERCA pumps act as additional calcium buffers, lowering the intrinsic release probability of vesicle release and increasing PPR. In addition, the presence of ER allows ongoing activity to trigger calcium influx from the presynaptic ER via ryanodine receptors (RyRs) and inositol trisphosphate receptors (IP3Rs). Intracellular stores and their associated machinery also allows a synapse with a low release probability to operate more reliably due to attenuation of calcium fluctuations. Finally, blocking ER activity in the presynaptic terminal mimics the pathological state of a low facilitating synapse characterized in animal models of Alzheimer’s disease, and underscores the critical role played by presynaptic stores in normal function.

## 1 Introduction

Synaptic transmission at small central synapses such as CA3-CA1 Schaffer collaterals is a tightly orchestrated event. Arrival of an action potential at a presynaptic bouton triggers voltage-dependent calcium channels (VDCCs) to initiate vesicle exocytosis. The spatial organization of each of the components involved in synaptic transmission (size of the docked pool of vesicles, arrangement of VDCCs, etc.) and their biophysical properties (concentration, binding rates and diffusion rates) decide the success of transmission (release probability of neurotransmitter) and play a critical role in constituting its plasticity profile^1^. At the CA3-CA1 synaptic terminal, the transmission success in response to an input at this synapse is conspicuously low, typically about 10–20%^2^. Instead, this apparent configuration flaw turns out to be an important feature that allows vesicle release probability to be tunable and the synapse to be highly plastic. The CA3 presynaptic terminal has a relatively small readily-releasable pool (RRP) size of 5–10 vesicles that are available for release. This synapse requires a delicate balance between conserving the vesicle resource and facilitating transmission capability^3^. A synaptic arrangement motif of a significant number of VDCCs albeit at a relatively large average distance from the site of vesicle release at the active zone (AZ) aids this facilitation^1,4^. The extended distance attenuates calcium concentration at the AZ and ensures a low baseline probability of vesicle release. The large number of VDCCs guarantees abundant bulk calcium that can modulate subsequent releases.

CA3 neurons have ER that is closely associated with the presynaptic terminal^5,6^. In reconstruction studies of rat CA3-CA1 synapses wherein around ~400 boutons were sampled, all boutons without exception had a substantial presence of ER^7,8^. ER acts as an internal calcium reservoir. Calcium ions are sequestered in the ER through the action of Smooth Endoplasmic Reticulum Calcium transport ATPases (SERCA) pumps and are released into the cytosol via either ryanodine receptors (RyRs) or inositol triphosphate receptors (IP3Rs) or both. Despite the ubiquitous presence of ER, its role in changing presynaptic calcium dynamics remains unexplored due to experimental difficulty in disambiguating the contributions of different sources of calcium.

Here, we investigate the contributions of ER to calcium signaling and short-term plasticity. We used a detailed computational model (see fig. 1) of the CA3 presynaptic terminal that incorporated essential components of neurotransmitter release, organized in a spatially realistic canonical bouton that allowed for *in silico* experiments. The main components that determine calcium dynamics in the presynaptic terminal are VDCCs, Plasma Membrane Calcium ATPases (PMCA), calcium buffers (predominantly calbindin in the presynaptic terminal), SERCA pumps, functional RyR and IP3R^9–14^. We simulate the 2D and 3D diffusion of individual molecules and their reactions and product formation using Monte Carlo algorithms in the designated synaptic spatial domain (details in Methods). We had previously shown that the model’s biophysical properties (diffusion constants, concentrations, kinetic schemes, binding rates) allow quantitative modeling of some characteristic transmitter release data at this synapse (time scales of release, initial release probabilities, amplitude of release etc.)^1,2, 15–17^

**Figure 1:**
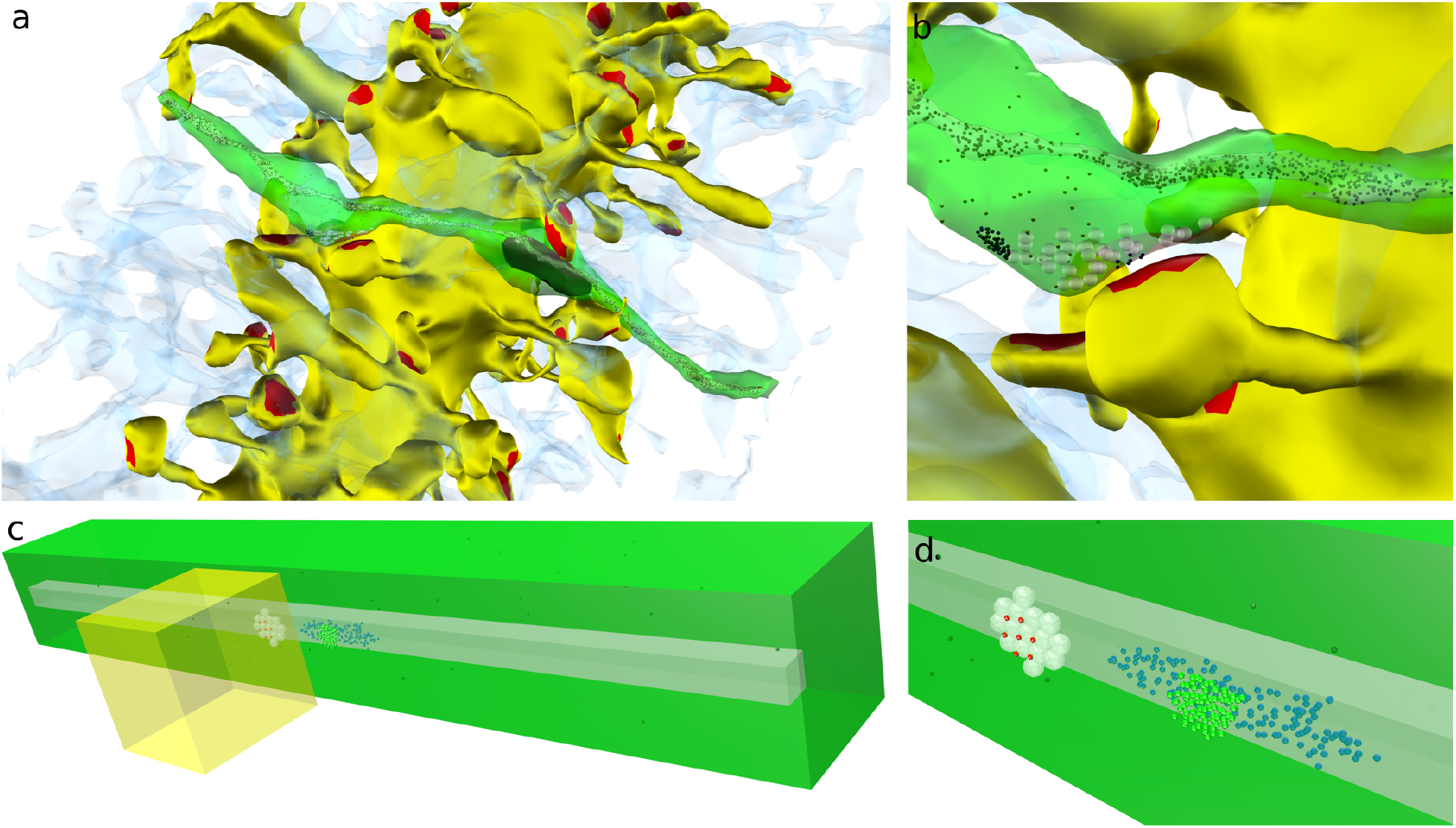
(a) Reconstruction of CA3-CA1 region in a rat hippocampus. Large yellow structure represents a CA1 dendrite with red patches being the postsynaptic densities. Axons of CA3 pyramidal neurons are coloured green (only one shown). ER inside the axon is shown in grey. Astrocytes are shown as semi-transparent blue structures. (b) An en passant synapse formed by CA3 axon onto CA1 dendrite. (c) Canonical model of a single CA3-CA1 synapse (same colour scheme used as for the reconstruction). (d) A closer look at the presynaptic terminal in the model shows docked vesicles (white), VGCC (green spheres) and RyR channels (blue spheres present on the ER).

One main advantage of our approach is we can evaluate in detail the effects of specific features of our presynaptic terminal. For example, Although presynaptic RyRs have been extensively reported^6,18^, the expression of group I mGluR (type 1 and 5) associated with IP3 production is controversial^19,20^. We can vary our control synapse to address the consequences of having or not having active IP3Rs. We can also investigate varying the baseline release probability, as suggested by Murthy et al.^17^. Finally, we can investigate the extent to which a detailed presynaptic reconstruction is critical by comparing results from our canonical geometry with those from more complex reconstructed morphologies (see fig. S2).

## 2 Results

### Synaptic transmission is a tightly choreographed event

Arrival of an action potential (AP) at an axonal bouton leads to the opening of VDCCs and a consequential large *Ca*^2+^ influx through the channels (fig. 2a). The rapid increase and fall in calcium concentration closely follows the voltage profile (fig. 2a). The PMCA pumps are vital in keeping calcium response brief in the bouton (fig. 2b). Additionally, the bulk calcium signal is governed by the mobile calcium buffer calbindin (fig. 2c) as it efficiently limits free calcium in the cytosol. However even as a large amount of calcium is extruded out, it takes ~100 ms for the synapse to recover to the 100 nM resting level calcium concentration. These lingering calcium ions are involved in facilitation of release at the next release pulse. Fig. 2d describes the local calcium concentration sensed by the docked vesicles at the AZ that leads to a characteristic release probability of ~20%. The peak bulk calcium concentration at this synapse (fig. 2d) is roughly one-half magnitude lower (at ~6 *μM*) than that of the calcium response at the active zone. The calcium sensors for vesicle release, Synaptotagmin I (synchronous release) and Synaptotagmin 7 (asynchronous release)^21^, can reach the threshold for release if an appropriate number of calcium ions diffuse to the AZ and bind to them. The distinctive release profile of transmitter release is a result of a low-affinity and fast calcium-binding site of Synaptotagmin I for immediate release during high calcium concentrations and a high-affinity but slow calcium binding property of Synaptotagmin 7 for late release. The resulting SERCA and calbindin activity as seen in fig. 2c in response to influx of calcium from VDCCs is fast enough to capture free calcium. Fig. 2e depicts the average opening of RyR and IP3R and the calcium flux through them in response to a single AP (Fig. 2a). The calcium released via these receptors are minor contributors at these short timescales; *Ca*^2+^ influx through VDCCs is much greater than through these receptors (see fig. 2a). Neurotransmission in response to a single AP arriving at the synapse averaged over 2000 trials is shown in fig. 2f. Fig. 2 describes how each of these presynaptic components can potentially change calcium signaling and synaptic transmission at the CA3-CA1 synapse.

**Figure 2:**
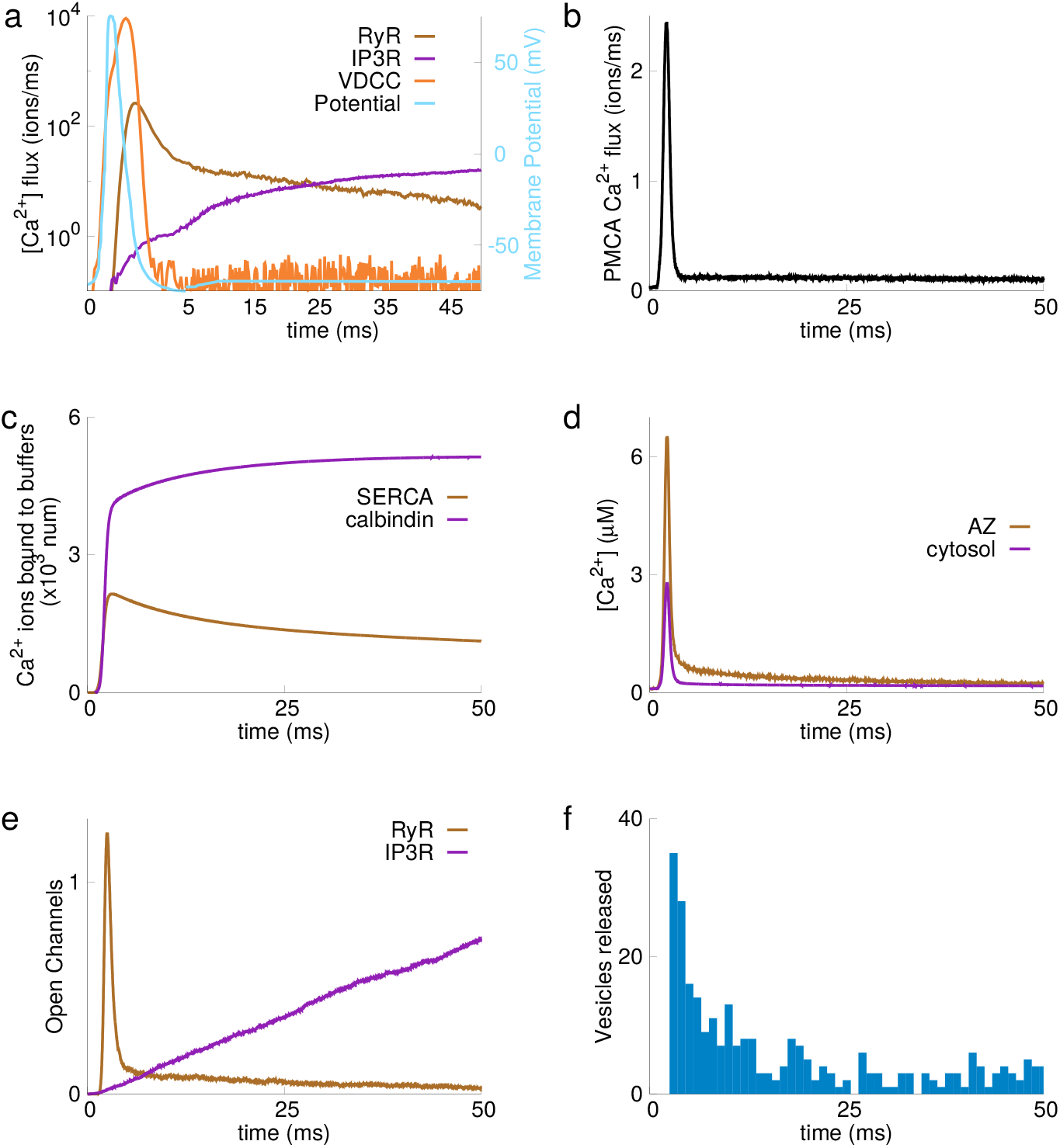
Behavior of different components of the model in response to a single initial AP during a time window of 50ms. (a) AP and *Ca*^2+^ Influx: Depolarization due to AP leads to opening of VDCC and subsequently, opening of RyR and IP3R, allowing influx of *Ca*^2+^ ions. *Ca*^2+^ ions bind RyRs when *Ca*^2+^ conc. increases in response to AP, opening the channel. RyRs respond fast and open and close in synchrony with *Ca*^2+^ conc. First 5 ms are stretched along the time axis for clarity. (b) PMCA Pumps: Actively removes *Ca*^2+^ from cytosol to maintain a base level *Ca*^2+^ conc. of 100 nm. (c) *Ca*^2+^ buffering: During an action potential (AP), large part of *Ca*^2+^ influx from voltage-dependent calcium channel (VDCC) are buffered by *Ca*^2+^ buffers (calbindin) and SERCA pumps. (d) *Ca*^2+^ concentration: *Ca*^2+^ influx from VDCC and RyRs, buffering by calbindin and SERCA pumps and pumping out by PMCA pumps leads to a characteristic *Ca*^2+^ signal in the presynaptic bouton. *Ca*^2+^ concentration is measured in the vicinity of active zone (marked as AZ) and global *Ca*^2+^ conc. is measured in the whole presynaptic bouton (marked as cytosol). AZ experiences higher concentration of *Ca*^2+^ than measured in the whole bouton and takes longer time to come back to its base-level conc. (e) Open Receptors: Ryanodine and IP3 receptors show distinct response to a single AP. RyRs open fast, almost in synchrony with the calcium influx while IP3Rs take much longer to open. (f) Vesicle Release: This characteristic rise in *Ca*^2+^ concentration profile leads to fusion of synaptic vesicles releasing neurotransmitter in synaptic cleft. Histogram of vesicle fusion event in 2,000 trials is shown in the figure.

### Blocking ER compromises short term plasticity

The neurotransmitter release probability of a synapse is thought to be an intrinsic property that arises out of the number of calcium channels, their spatial arrangement with respect to the active zone and the size of the RRP. Most small hippocampal synapses operate in the regime of low release probability^2^. Synapses with larger numbers of calcium channels have a higher intrinsic release probability Pr (fig. 3a); however, the dependence of Pr on calcium channels becomes steeper when SERCAs are blocked. A control synapse with ER will always have lower Pr compared to a synapse without ER (for example, with 90 VDCCs Pr = 0.2 (with ER) and Pr = 0.4 (no ER)). This effect is due to SERCA pumps, not to the ER release channels.

**Figure 3:**
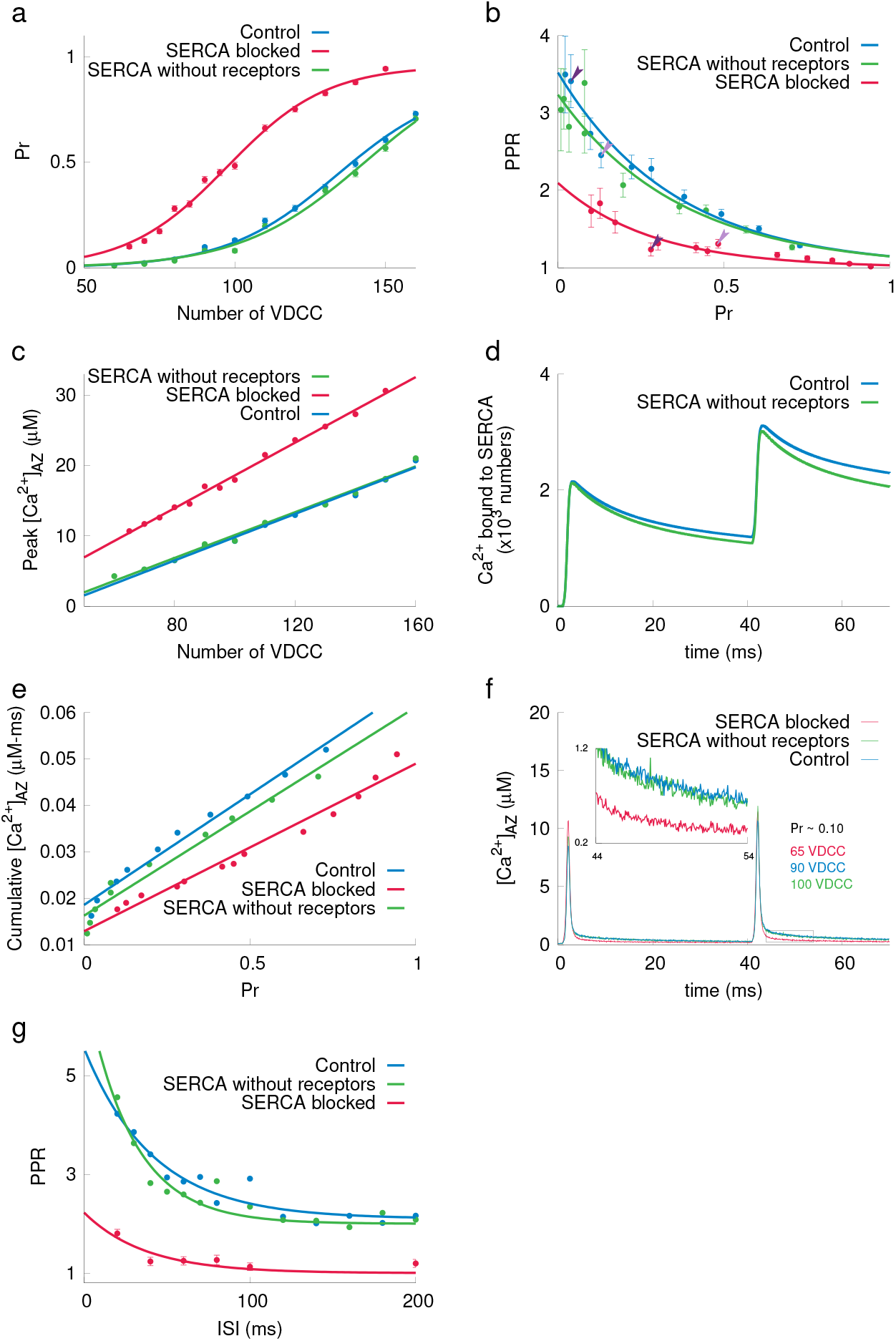
ER’s contribution to PPR (a) Variation of Pr with number of VDCCs. When SERCAs are blocked (red), Pr is drastically higher than in Control (green) and when only ER channels are blocked (blue). (b) Inverse relation of paired pulse ratio and intrinsic release probability for various synaptic configurations. Control synapses with ER and receptors, SERCA blocked and only SERCA and no receptor. The arrows indicate the PPR corresponding to the same number of VDCCs for Control and SERCA blocked. (c) Variation in peak 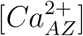 with number of VDCCs. When SERCAs are blocked, [*Ca*^2+^]_*AZ*_ is higher than in control synapses (ER receptors blocked shows similar profile as control). (d) Amount of *Ca*^2+^ ions bound with SERCA in response to 40 ms ISI paired pulse protocol. (e) Cumulative calcium concentrations at AZ for an interval of 20ms after initiation of second AP is plotted for various Pr. (f) Local calcium concentration at the AZ is shown for Pr = 0.10 Inset (zoom into box): details of base level 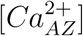 concentration after second AP. (g) Variation of paired pulse ratio for different ISI.

To study plasticity, we first use the paired-pulse ratio, a classic measure of presynaptic STP to characterize plasticity profiles. As in classic studies of small hippocampal synapses, PPR is defined as ratio of probability of neurotransmitter release for the second stimulus (*Pr*_2_) divided by the probability of neurotransmitter release for the first stimulus (*Pr*_1_), averaged over multiple trials^2,22^. The inverse relationship observed between the release probability and the PPR is a universal feature for small hippocampal synapses^2,17, 22^. One reason for this is that the release probability is bounded by 1, and at a high initial release probability, there is a limit to how much more it can be increased, thus limiting the size of the PPR. For synapses with very high intrinsic release probabilities, depletion of the small RRP (due to multiple vesicles being released via the first stimulus) overwhelms calcium-driven facilitation and gives rise to low PPR. At the typical operating point of low release, synapses can have a large PPR range, tunable in an activity-dependent manner^23^. Fig. 3b shows PPR responses (with a 40 ms ISI) to a complete range of intrinsic release probabilities (max PPR ~6). Blocking ER activity (SERCA blocked) substantially diminishes the PPR (fig. 3b, red curve) at a fixed release probability. This is especially true in the low release probability regime where most CA3-CA1 synapses function.

We now describe the mechanism whereby synapses with ER can sustain higher PPR at low Pr. SERCA pumps have a high affinity and fast binding for calcium^24^. As a consequence, the peak amplitude of calcium signals measured at the active zone for a particular synaptic configuration is lower than it would be without SERCA pumps (fig. 3c). This enables rapid sequestration of calcium into the ER (fig. 3d). This fast buffering action of SERCA magnifies the storage capacity of calcium at the synapse and hence gives rise to a smaller release probability compared to the same synapse without ER. But eventually the calcium unbinds from the SERCA: as a result, a larger cumulative and a shorter peak calcium amplitude occurs in synapses with ER compared to those without ER over a wide range of intrinsic release probabilities (fig. 3e and f). Simulations for a shorter 20 ms ISI is shown in supplementary fig S3. The longer lasting calcium transient contributes to facilitated vesicle release in synapses with ER. This facilitation does not have to compete with vesicle depletion if the baseline release probability is small. The enhanced PPR for synapse with ER occurs over a wide range of ISIs (fig. 3g). These results reveal an important, previously unaccounted role for SERCA in enhancing short-term plasticity. Note that the binding rates and affinities used here correspond to an ER refilling timescale as seen in experiments^25,26^. However, these results are not sensitive to reasonable variations in SERCA densities and rates of SERCA binding (see fig. S6). Calcium release via ER channel opening is not a major contributor for time scales relevant to PPR.

### Contribution of intracellular stores to facilitation in response to a stimulus train

Next, we investigated the effects of ER on synapses stimulated with a train of APs. Fig. 4 describes synaptic activity driven by 20 AP arriving at the rate of 20 Hz (see supplementary material for stimuli at 50 Hz and 10 Hz, fig. S4 and S5, respectively). The *in silico* experiments were conducted for four different synaptic configurations 1) Control (RyRs + SERCA), 2) IP3R (with SERCA), 3) SERCA without receptors and 4) SERCA blocked (no ER). All four configurations of synapses show a peak in facilitation (fig. 4a) defined as ratio of vesicle release probability of the *n^th^* stimulus to the first stimulus. The initial positive slope in facilitation is apparent when plotting the probability of vesicle release as a function of stimulus number (fig. 4b). As before, in PPR simulations, a high baseline release probability is linked to low facilitation (fig. 4, compare a and b). The increase in vesicle release rate as the train progresses leads to a rapid depletion of the RRP (fig. 4i). This competition between availability of vesicle resources (RRP size = 7) and increase in release rate lowers facilitation after the initial increase for all synaptic configurations. The vesicle recovery time at a single site is 2.7 sec^2^).

**Figure 4:**
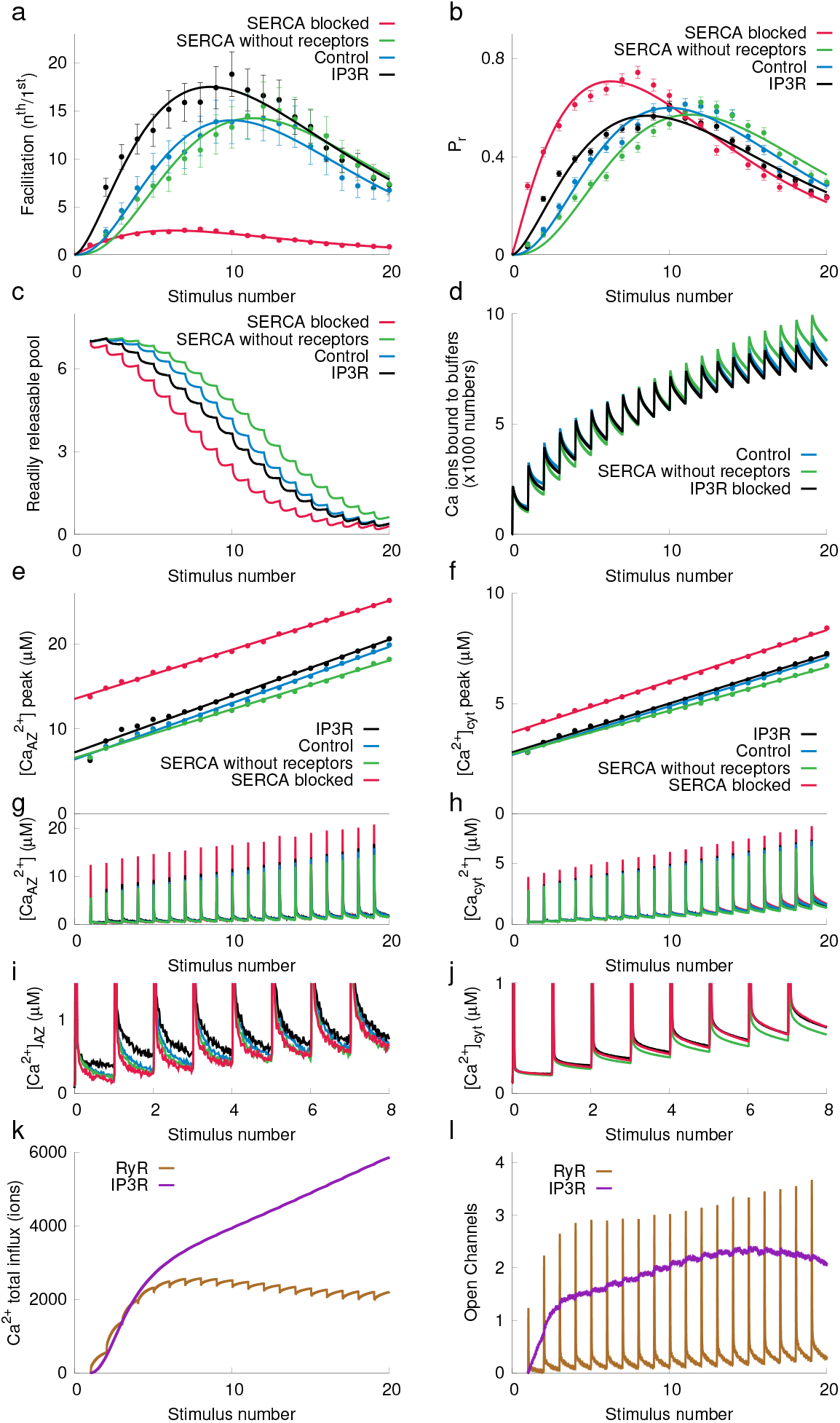
Response to 20 pulses at the rate of 20 Hz. (a) Frequency facilitation for a synapse with SERCA blocked (red) is lowest compared to a range of synaptic configurations that include ER (green, blue and black) (b) Release probability for a synapse with stores (red) and synapse with ER (green, blue and black). (c) Corresponding change in vesicles available for release (RRP). (d) Calcium activity at SERCA pumps. As calcium peaks rise with successive input pulses (e, f, g, h, i and j), release probabilities rise. This rise in release probabilities is suppressed by vesicle depletion (c). Peak [*Ca*^2+^]_*AZ*_ is shown (e) along with complete [*Ca*^2+^]_*AZ*_ profile (g) and its base level values (i). Peak calcium concentration is drastically higher when SERCAs are blocked. (f, h, j). (k) [*Ca*^2+^] flux from RyR and IP3R is shown. IP3Rs are slow low but sustained over a longer period of time compared to RyRs. (l) Corresponding open RyR and IP3R channels.

We can analyze this process in more detail. As in previous protocols, SERCAs rapidly bind a large amount of incoming calcium ions during an AP, thus decreasing the calcium ions available for binding to the synaptotagmins compared to when SERCAs are blocked (fig. 4j). The probability of vesicle release is primarily dictated by peak calcium concentration. Since the peak of the calcium response is maximum when SERCAs are blocked (fig. 4c, d, e and f), the highest vesicle release rates and lower facilitation are associated with this configuration (fig. 4a, b). In the presence of ER, peak values of calcium concentration are lower and basal (post AP) calcium concentrations are higher because of calcium release from RyRs and IP3Rs (fig. 4i and j), further enhancing facilitation. In order to open and release calcium, both of these receptors have to bind calcium. A slower calcium binding rate and the additional requirement of IP3 binding renders IP3Rs slower. However, a slow closing time and high affinity for calcium allows IP3Rs to sustain calcium release for several seconds, long after the VDCC have closed. Owing to these biophysical properties, the contribution of IP3 receptors to basal calcium is significantly higher than that of the RyRs (fig. 4 k and i). RyRs, on the other hand, bind calcium faster but have a lower affinity, requiring higher calcium concentrations to become activated (fig. 2a). Therefore, for a train of stimuli, IP3Rs contribute to facilitation in addition to SERCA.

### Increased reliability mediated by stores machinery

Experimental data on spatial navigation and sensory processing^27,28^ suggests that information is encoded in small hippocampal synapses via a finely tuned activity rate. This is achieved despite the highly stochastic nature of individual synapses operating in the low release probability range. Here, we show that intracellular calcium stores and their associated machinery may contribute to the reliability of synapses with low release probabilities.

We focus on the conditional probability, *P*_11_, that a successful transmission event in response to the first stimulus is followed by another successful transmission event in response to the second stimulus. A graph of this probability is shown in figure 5a. *P*_11_ is higher for synapses with calcium stores across all *Pr*_1_. This suggests that a synapse with calcium stores predicts subsequent activity more reliably. This is a consequence of the longer lasting, higher amplitude calcium signals with lower fluctuations in a synapse with calcium stores. At fixed *Pr*_1_, a larger number of VDCCs are needed in control synapses, and hence, the coefficient of variation (CV) of the calcium signal is lower (fig. 5b). Note that SERCAs keep up with demands of high calcium at high Pr. Correlation between maximal % of SERCA binding sites during a stimulus and peak calcium during a stimulus for our canonical synapse configuration is shown in figure 5c. Figure 5d describes decay of CV of peak calcium concentration with SERCA occupancy. CV of calcium concentration was higher with SERCA blocked (fig. 5e). These data describe how buffering by SERCA and higher level of sustained calcium after the first pulse in control synapses further lowers calcium fluctuations. The less noisy calcium signal associated with control synapse may also have implications in other calcium dependent downstream signaling including vesicle recycling^29^. In summary, intracellular calcium stores allow synapses to operate at low intrinsic release probability and yet exhibit large, reliable facilitated release caused by longer calcium driven synaptic traces.

**Figure 5:**
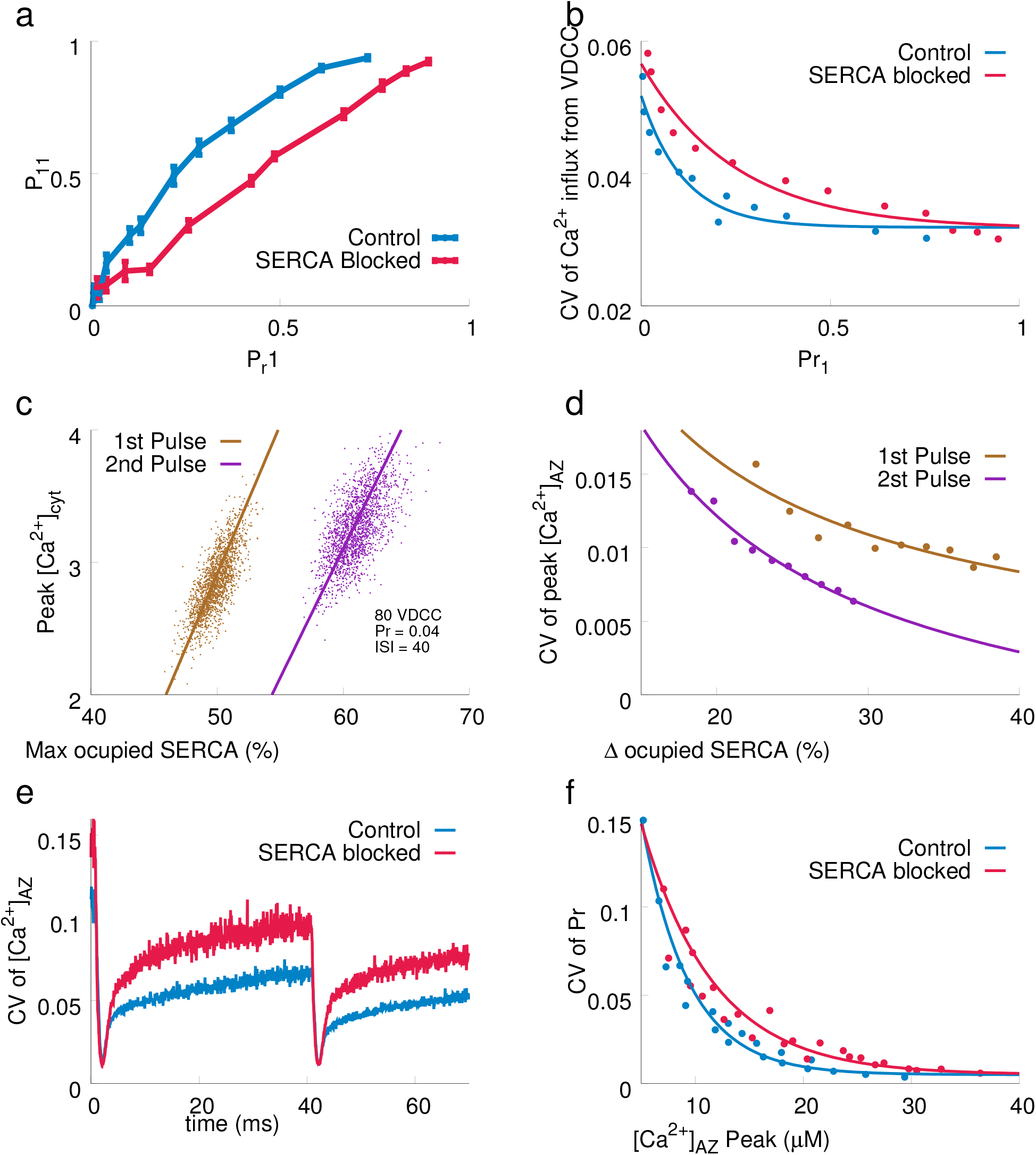
(a) Probability of vesicle release after the second AP given that the first AP results in a successful vesicle release (*P*_11_) as a function of *P r*_1_. (b) CV of calcium flux through VDCCs as a function of *P r*_1_. (c) Correlation between peak cytosolic calcium concentration and the maximum occupancy of SERCA pumps for (1000 trials) for a release probability ~0.2. (d) Variation in CV of peak [*Ca*^2+^]_*AZ*_ as a function of occupied SERCA sites. (e) CV of [*Ca*^2+^]_*AZ*_ for a synaptic configuration with 80 VDCCs. (f) CV of *P r*_1_ varying with peak [*Ca*^2+^]_*AZ*_.

### Intracellular calcium store necessary for PPR requirement of a CA3-CA1 synapse

Thus far we have described the mechanisms by which SERCA and other components associated with calcium release from ER influence synaptic transmission. Here we compare the predictions of our model with experimental observations of short-term plasticity at CA1-CA3 synapses^2,17^.

In figure 6a, we show PPR for the entire range of intrinsic release probabilities as predicted by the model (control and with SERCA blocked) and as measured in two independent experiments. Simulations of control synapses that include presynaptic calcium stores are in better agreement with experimental data for both 40 and 50 ms ISI. Experimental data^2,17^ and simulation data were fit to

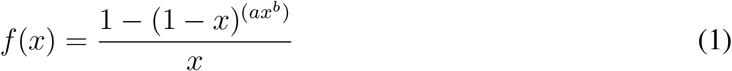

**Figure 6:**
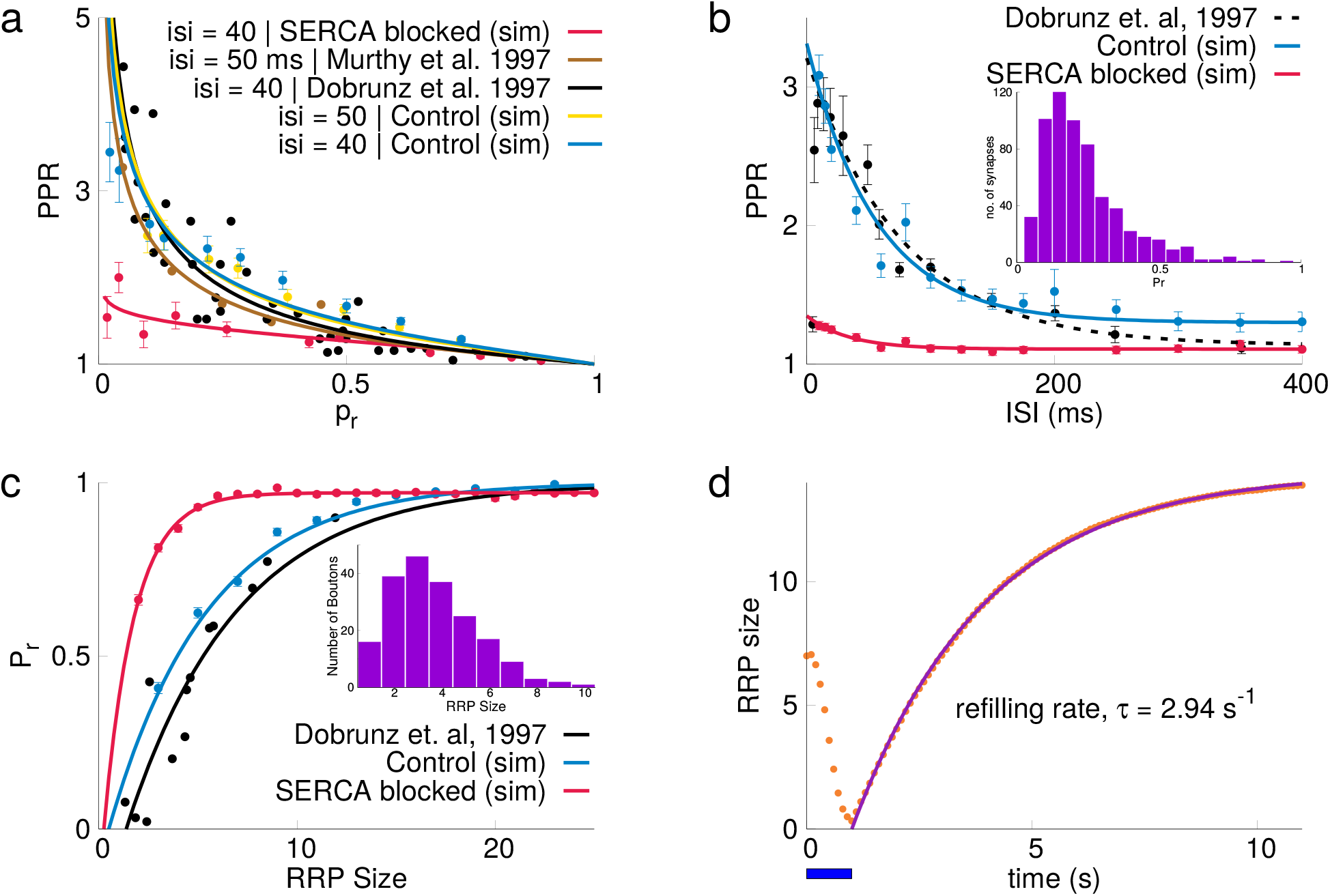
Comparison of simulation results with experimental data. (a) Inverse relation of PPR with Pr for ISI 40 ms and 50 ms (b) PPR also reduces with increasing ISI for both synapses with and without stores. Inset shows the distribution of Pr for the set of synapses used in the experimental result and simulations(redrawn from Dobrunz et. al, 1997. (c) Variation of Pr with RRP size (experimental data: Dobrunz et. al, 1997). Distribution of RRP size used for simulations is shown in inset (redrawn from Dobrunz et. al, 1997). (d) RRP was depleted in by a 20 hz stimulus for 1 sec (indicated by horizontal blue bar) and vesicle refilling was observe and fitted to an exponential curve shown as blue curve (refilling rate, *τ* = 2.94*s^−1^*).

Eq. 1 is a good fit for our simulation data only when the ER and the release mechanisms for intra-cellular stores were included (fig. 6a). Fig. 6b shows PPR as function of ISI. A part of the calcium signal in response to the first AP may enhance calcium response to the second AP, modulating release and PPR. As the ISI increases, the probability of influencing the second release by residual calcium decreases. Accordingly, the PPR decreases with increasing ISI in all scenarios. In experiments by Dobrunz et al.^16^, PPR as a function of ISI was calculated for ensembles of CA3-CA1 synapses with a distribution of release probabilities described in Murthy et al.^17^. In order to replicate these experimental protocols, we incorporated the same distribution of release probabilities in our model and measured the PPR for the same range of ISI. Without making any further changes to the model or the model parameters, the PPR response of the model closely followed the experimental measurements only in control synapse when the stores were included. In contrast, when the stores were blocked (SERCA blocked), the PPR response was reduced drastically and was not in agreement with experimental data.

The dependence of release probability on RRP size was measured in a different set of experiments by Dobrunz et al.^2^. These experiments were carried out with extracellular *Ca*^2+^ concentration at 4 mM, roughly double the physiological value; hence for an accurate comparison, our model calculations included this doubling of extracellular calcium concentration for both the control synapse and the synapse with calcium stores blocked (see fig. 6c). A clear match with experiments was again found with the control synapse when intracellular stores were included. When stores were blocked, we found a much higher vesicle release probability, which saturated to almost 1 for a very small number of vesicles (~5). This higher *Pr*_1_ response in simulations with calcium stores blocked can again be attributed to absence of buffering by SERCA. The timescales for vesicle recycling can critically modify both PPR and release probabilities. The vesicle recovery after depletion of our model shown in fig. 6d matched experimental observations.

## 3 Discussion

Our simulations predict that presynaptic ER in CA3-CA1 hippocampal synapses contributes to the short-term plasticity profile. The ER-mediated increase in facilitation that we have simulated explains experimental observations. Recent studies have demonstrated that in CA3-CA1 and other synapses, facilitation is lowered when ER is blocked (with thapsigargin), over a range of ISIs and stimulus frequencies^30,31^. Interestingly, Zhang et al.^31^ also show that this effect of lowered facilitation quantitatively mimics Alzheimer’s disease (AD) synapses (synapses from a presenilin model of AD^30^). These data suggest ER modulation of plasticity is critical for normal function and disrupted ER signaling is a potential locus for the pathogenesis of AD. Our model outlines the distinct mechanisms by which SERCA and related intracellular mechanisms differentially modulate short-term plasticity over a range of time scales.

The presence of ER is reported in both axonal and dendritic compartments^6,32^. Only about 20% of spines have ER and it is over-represented in the larger, stronger synapses^33^. In contrast, the ER in CA3 axons extends through all boutons^7^. Amongst its other functional roles, the endoplasmic reticulum can directly modulate downstream calcium signaling. Here, we report on two distinct mechanisms of ER modulation of synaptic plasticity via its action on calcium. Over shorter time scales of few tens of milliseconds, SERCA pumps, via their fast calcium binding sites, rapidly bind incoming calcium from VDCCs. This lowers the free cytosolic calcium leading to lowering of release probability of vesicles. Our control synapses that include ER display an the inverse relationship between PPR and release probability of vesicle releases at the CA3 presynaptic terminal, a well-established experimental relationship^2,34^ (see fig. 6a). Fast action and high affinity binding are characteristic features of SERCA^24^ that are essential for physiological refilling of the ER. These biophysical properties of SERCA make it highly effective at lowering bulk calcium. SERCA’s calcium buffering properties in turn cause a lower *P r*_1_. A low intrinsic Pr allows a synapse to operate with a higher flexibility in short-term plasticity. The reported time-scale of calcium transient decay and the ER refilling rate constrain the binding rates and expression levels of SERCA^25,26^.

In summary, the presence of ER allows synapses to capture a greater amount of VDCC calcium via SERCA binding, keeping the free calcium low and the consequential intrinsic release probability low. Upon a second stimulus calcium is already bound to SERCA, resulting in increased free calcium, increased release probability and hence PPR>1. Furthermore, a low release probability ensures high PPR for control synapses compared with synapses with no ER. The dramatic shift in PR is shown in fig. 3a for a synapse with identical intrinsic properties except presence of ER (illustrative values indicated by arrows). The actual contribution of calcium release from RyR and IP3R over these shorter time scales is minor. Over longer time scales, for example in response to a sustained train stimulus, calcium release from the ER due to opening of RyR and IP3Rs can amplify VDCC calcium entry and also modulate facilitation.

At the CA3 terminal there is a trade-off between a wide range of short-term plasticity and reliability. However, spatial location and sensory information is encoded in the firing rates of pyramidal cells in the hippocampus. Changes in sensory stimuli such as odor or color also lead to changes in the firing rates of these cells, called rate remapping. Lesions to the hippocampus that impair rate remapping can compromise sensory discrimination^27^. Reliable firing rates could be maintained despite the stochasticity of vesicle release if a large number of synapses are activated with bursts of spikes rather than isolated spikes to recruit facilitation. However, it has also been argued that the hippocampal rate code describing a location in an environment is represented by activity of a small number of CA1 pyramidal neurons (population rate code)^35,36^. The alternative strategy would be to operate at high vesicular release probability, but at high Pr, the small RRP size at this synapse would steer the synapse towards much quicker depletion. The ER allows synapses to have low intrinsic release probability but still exhibit relatively higher vesicle release predictability.

Both IP3R and RyR expression have been reported in presynaptic terminals; however, the presence of presynaptic RyR is more prevalent^6, 11, 18, 30, 31, 37, 38^. Binding of calcium alone triggers opening of ryanodine receptors, leading to calcium release. Conversely, the opening of IP3 receptors requires both IP3 and calcium to be bound. Group I mGluR (Type I and V) associated with IP3 production have been reported in the hippocampus^39–41^. Spillover of glutamate released from vesicles can lead to IP3 production via mGluR and G-protein pathways (see methods for details) and lead to IP3R opening and consequential calcium release. Ryanodine receptors have high affinity and require a large calcium transient to open and hence directly follow the VDCC *Ca*^2+^ flux. IP3 receptors have low affinity and remain open for several seconds past the VDCC flux termination (fig. 4k,l). The distinct biophysical properties for each of these receptors provide a wide spatiotemporal range for calcium signaling that can be sustained for seconds and participate in several different forms of plasticity. Given the differential expression of these receptors, we have systematically investigated various synaptic configurations and the combined effect of each of these receptors as well as when they act independently^31^. We conclude that the presence of presynaptic ER allows a repertoire of multiple time scales for calcium signaling, and for other downstream molecular signaling mediated by ER dynamics^42,43^.

The model parameters in our canonical synapse have been chosen to account for the observed release timescales, release peaks, calcium affinities and effective calcium diffusion^15^. The parameters of the intracellular calcium release machinery (SERCA Pumps, RyR receptors) have been calibrated by the steady-state concentration of calcium in the ER and cytosol and by the refilling rates of the ER. The vesicle release rate and PPR profile was robust to realistic variation of these parameters and synaptic geometries (see supplementary information). The relationship between PPR and initial release probability has been quantified by several laboratories^2,17^. Our strategy has been to use a well-constrained biophysically detailed model to predict the functional implications of intracellular stores. Physiological measurements of paired-pulse facilitation and release probability is governed by several variables, including the resting calcium level, the peak calcium flux through VDCCs, RRP sizes, the calcium sensor for vesicle release, calcium buffering and the inter-stimulus interval. Each of these variables has a nonlinear relationship with the local calcium signal as seen by the vesicles that ultimately affects release. A biophysical model that quantitatively agrees with measurement of the PPR for different protocols should have physiologically realistic implications. The canonical model that includes intracellular calcium stores, without adjusting for any parameter, is in good agreement with several independent lines of investigations on the CA3 presynaptic terminal.

The comparisons presented strengthens the essential contribution of presynaptic intracellular stores towards the intrinsic plasticity of the synapse. The model described here is high dimensional and is capable of describing several synaptic mechanisms characteristic of a CA3-CA1 synapse. Additionally, we also simulated the same PPR protocol and train stimulus with appropriate model components (20 ms ISI) in 1) a specific instantiation of a CA3 axon from a 6×6×5 *μm*^3^ cube of neuropil from stratum radiatum of the hippocampus reconstructed from electron micrographs of 100 serial sections (See methods); 2) a synapse with simplified cuboidal geometry of same dimension as the reconstruction (see fig. S2). The robustness of the findings ensures that our results are general and not restricted to a specific geometry. A low Pr synapse not only allows for vesicle release to be tuned over a wider range but also allows a synapse to minimize energy use^44^.

The functional implications of various kinds of short-term plasticity remain enduring questions in neuroscience. Short-term plasticity modulates the functional efficacy of synaptic transmission at a millisecond-to-seconds timescale. In the mammalian brain, short-term synaptic plasticity influences the information processing function of synapses, enabling them to optimize network level computation^45^. Synapses with a low initial probability of release function as high-pass filters, while synapses with a high initial probability of release function as low-pass filters^46–48^. Abnormal STP is among the earliest indications of various forms of dementia^49^. We would need to extend the model to investigate contribution of ER to presynaptic LTP and the long-term costs of loss in STP as seen in AD. However, studies that show that low Pr (and high PPR) in CA3 bouton corresponds to higher magnitude of LTP directly connects STP to LTP^50^.

Several studies^51^ have shown that a biological system can achieve a physiological goal via multiple pathways. Because more than one set of parameters can model an experimental observation^52^, it would be imprudent to pin down a unique set. However validating the model on diverse experimental data restricts this parameter space substantially. But it is also true that despite this degeneracy in parameter space and redundancy of signaling pathways, a small aberration in these components can lead to pathological states. Strong validation for our conclusion that presynaptic ER has a critical functional role is based the presence of ER in all CA3 axons and that blocking ER lowers facilitation and mimics short-term plasticity profile of an AD axon.

## 4 Methods

MCell, version 3, a Monte Carlo Cell simulator, was used to carry out the simulations. Monte Carlo algorithms are used by MCell to simulate diffusion of individual molecules present either on a surface or in a confined volume. MCell carries out user-specified molecular reactions stochastically. We simulated an en passant axon segment with physiologic spatial distributions and concentrations of molecules. These simulations track each molecule and the relevant reactions to calculate spatiotemporal trajectories. Simulations were performed on a cluster with 1464 processing units dedicated for the neurobiology group in IISER, Pune. Several thousand trajectories (1000–5000) were simulated to arrive at an average trajectory.

### Model Components and Geometry

Simulations were carried out in 1) a presynaptic terminal with canonical dimensions (0.5 × 0.5 × 4*μm*^3^ volume, surface area: 8.5*μm*^2^)^15^ representative of a canonical CA3 presynaptic terminal with simplified cuboidal geometry, in 2) a CA3 axon reconstructed from EM data with volume 0.39*μm*^3^ (see fig. 1). The details of reconstruction of neuropil can be found on page 4 in Bartol et. al^8^, and in 3) a presynaptic terminal with simplified cuboidal dimensions similar to reconstructed synapse from EM images.

The canonical model is composed of three major geometrical components: cuboidal presynaptic and postsynaptic terminal and U-shaped astrocyte covering the synapse. Presynaptic terminal contains cuboidal ER compartment. Dimensions of these components are given in fig. 7. Molecular components of the model, their quantity and placement are listed in table 1. Reaction rates for the reactions are given in table 4 and model parameters are listed in table 2. Kinetic scheme for RyR, IP3R and SERCA pumps are provided in fig. 8, respectively, and that of calcium sensors is provided in Nadkarni et al. 2010^15^.

**Figure 7:**
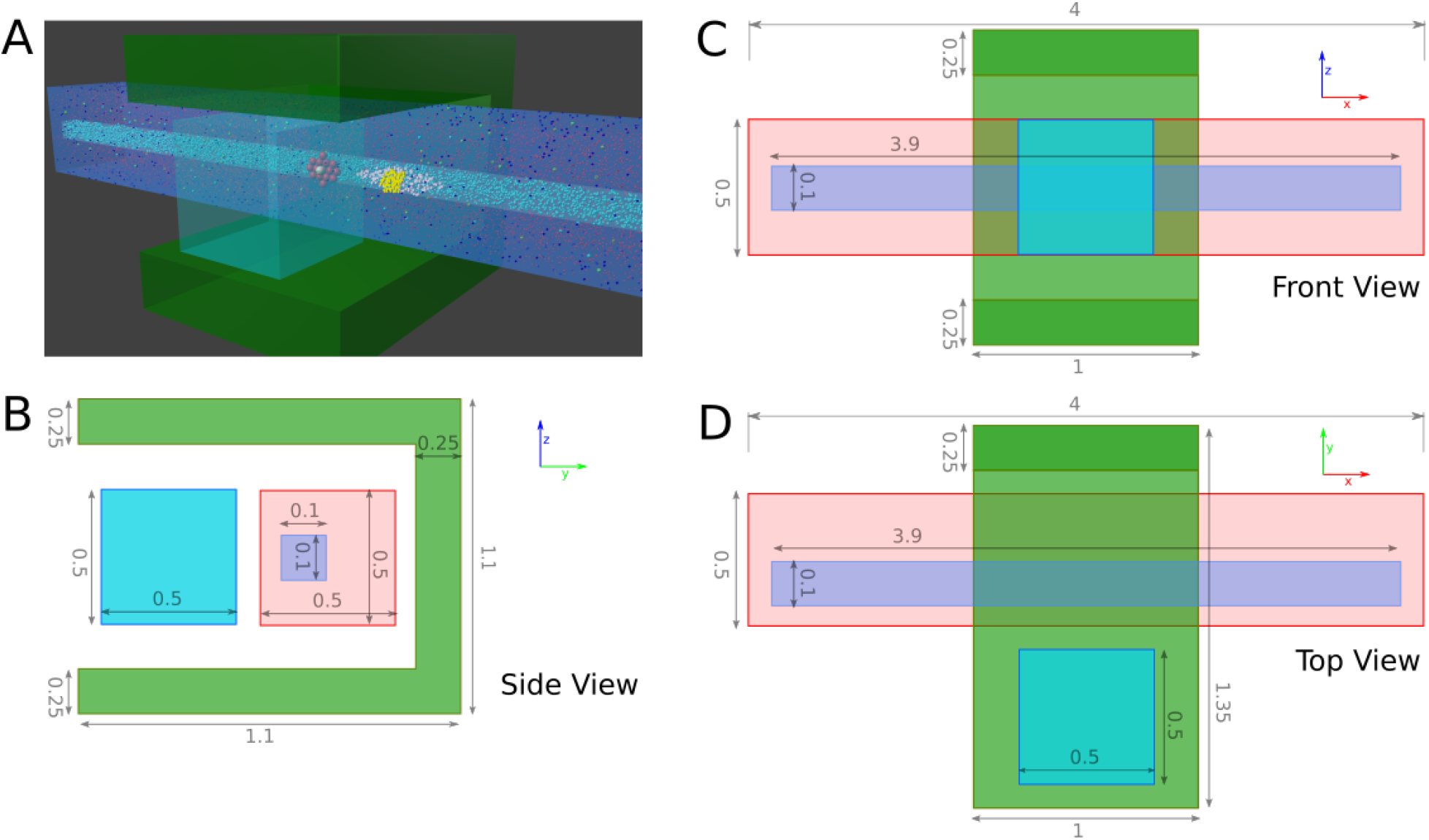
Geometry of the model. (a) Placement of important molecular species in the model. (b, c, d) Dimensions of the model (in *μm*) are provided in orthographic projection.

**Figure 8:**
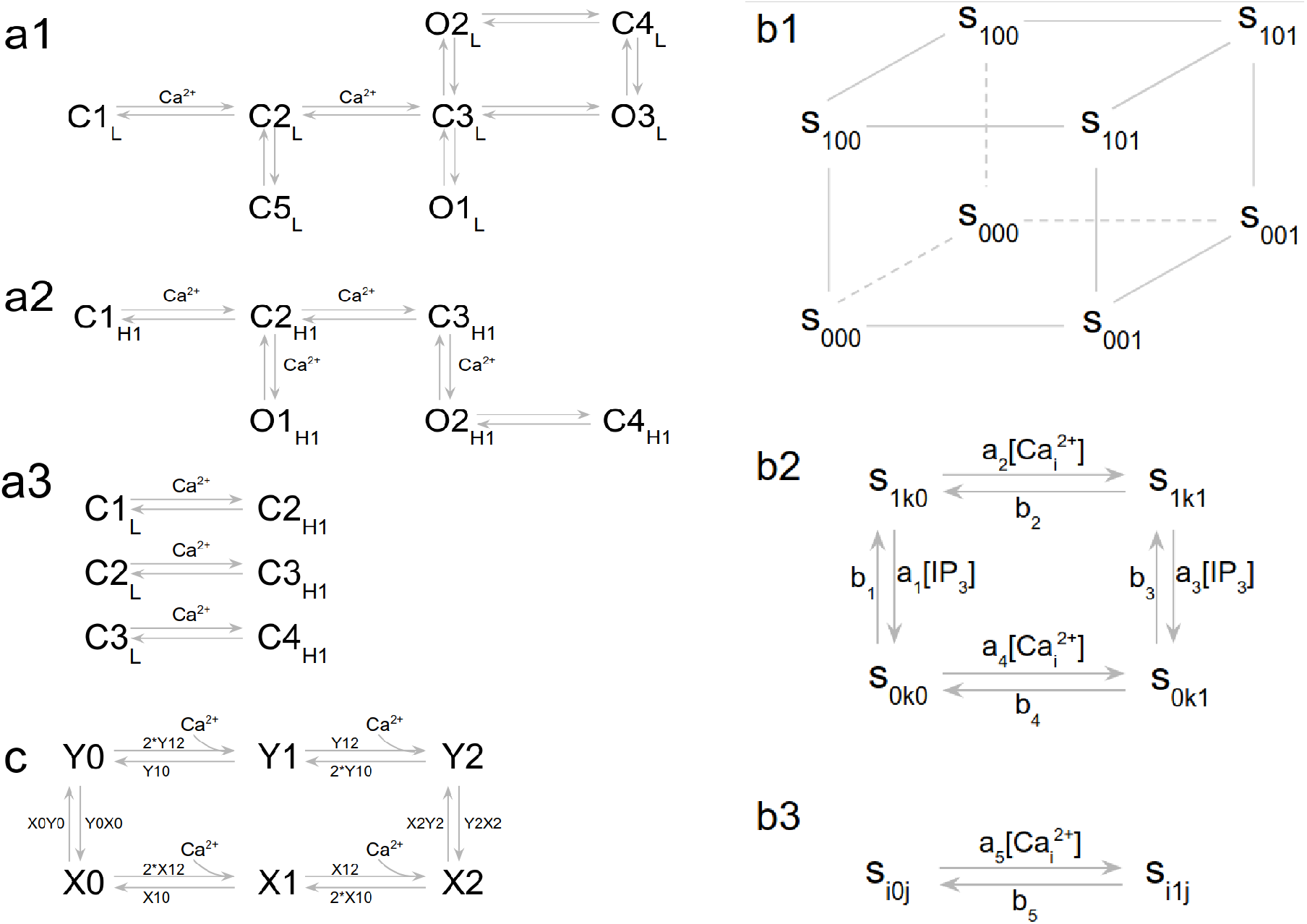
(a1–3) Kinetic Scheme for RyR-L mode, RyR-H1 mode, transition between L and H1 modes of RyR. (b1–3) Kinetic scheme of de young keizer model for IP3 receptors (adapted from Young et al.^53^) (c) Kinetic Scheme for SERCA pump.

**Table 1:** Components of the canonical model used for simulation.

| **Component** | **Quantity** | **Placement (measured from AZ)** |
| --- | --- | --- |
| RyR | 150 | 350 nm |
| IP3R | 50 or 0 | 150 nm |
| VDCC | 40 - 160 | 350 nm |
| Active Zone | 1 | 0 nm |
| PMCA Pumps | 180 | plasma membrane, uniformly distributed |
| Calcium Buffers | 45e-6 M | cytosol |
| SERCA Pumps | 15e-6 M | cytosol |
| Glu Transporters | 1000 per $\mu m^2$ | Glia surface, uniformly distributed |
| IP3 | 160e-9 M | cytosol |
| mGluR | 1000 | plasma membrane, uniformly distributed |
| $C a_{ER}$ | 250e-6 M | ER |
| $C a_{cyt}$ | 100e-9 M | cytosol |

**Table 2:** Parameters of the canonical model used for simulation.

|  |  |
| --- | --- |
| Calcium diffusion Constant (DCa) | $220 \text{ mm}^2/s$ |
| Calbindin diffusion constant (Dcb) | $0, 28 \text{ mm}^2/s$ |
| PMCA diffusion Constant (DPMCA) | 0 |
| VDCC diffusion constant (Dvdcc) | 0 |
| Glutamate diffusion constant (Dglu) | $200 \text{ mm}^2/s$ |
| IP3 diffusion constant (Dip3) | $2.8\text{e-}6 \text{ mm}^2/s$ |

**Table 3:** The different cases in which simulations are run.

| **Case** | **ER components present** |
| --- | --- |
| Control | SERCA Pumps and RyR |
| SERCA Blocked | none |
| SERCA without receptors | SERCA Pumps |
| IP3R | SERCA Pumps and IP3R |

**Table 4:**
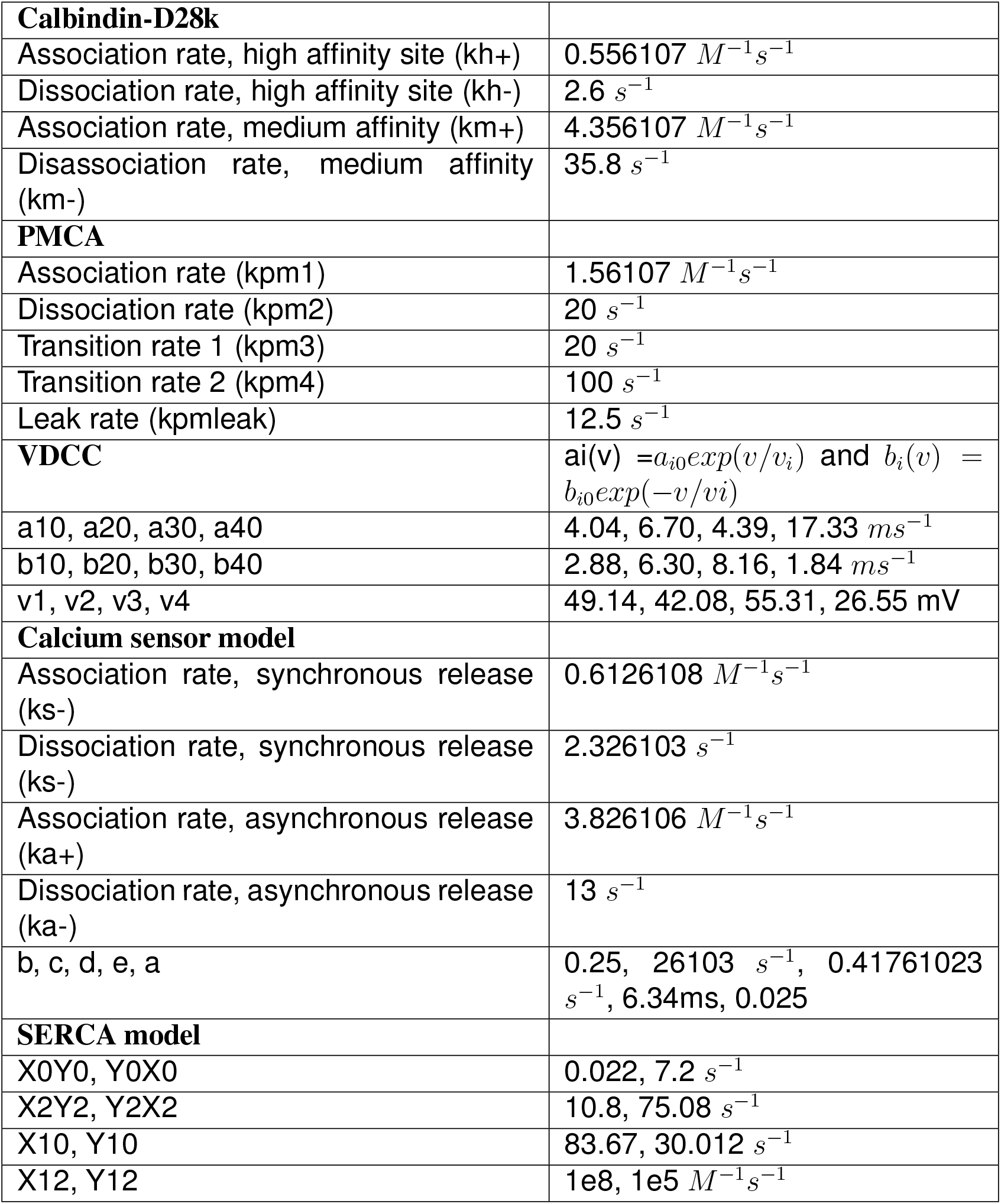
Reaction rates for the different kinetic schemes used in the simulation.

### Model Configurations

Model is setup to have base level Ca^2+^ concentration in cytosol at 100 nM and and ER Ca^2+^ concentration at 250 uM when there is no activity. Simulations have been performed in 4 different setups as mentioned in table 3. For frequency facilitation, simulations were run having 20 APs at 5, 10, 20 and 50 hz. For Paired pulse ratio, ISI was varied from 20 to 200 ms with an interval of 10 ms until 100 ms and then with a step of 20 ms until 200 ms. To vary the vesicle release probability, calcium influx through VDCCs was varied by changing the number of VDCCs from 40 to 160. Spatial calcium profile remains the same whether mobile or immobile calcium buffers are used (see fig. S1). We have therefore set calbindin diffusion to zero without loss of accuracy for computational efficiency. Initial RRP size has been kept 7 throughout all the simulations unless otherwise mentioned. The activity of SERCA pumps in response to calcium influx on the ER membrane along with PMCA pumps on the cytoplasmic membrane maintain 3 distinct pools of calcium (resting calcium concentration of ~250 *μM* in the ER and 100 nM in the cytosol and 2 mM extracellular space). IP3Rs require binding of both calcium and IP3 to get activated. Calcium binding to the inactivation site deactivates these receptors at high calcium concentrations^53^. Typically a feedback cascade initiated by glutamate released through vesicles and binding to presynaptic metabotropic glutamate receptors (mGluR) results in production of inositol trisphosphate (IP3) in cytosol^54^. The signaling cascade results in further opening of IP3Rs requires several seconds.

### Simulations

In simulations with a paired-pulse stimulus, the number of trials are: 5000 trials for VDCC=40–60, 2000 for VDCC=70–90, 1000 trials for VDCC=100–160. Release probability in response to an AP is calculated by counting the number of vesicles released during a period of 20 ms starting from the initiation of AP in all the trials. When more than one vesicles are released in response to an AP, it has been counted as one. This is in accordance with the definition of vesicle release probability: probability that at least one vesicle is released. Error in release probability was calculated using 1000 resampling with replacement and then mean and standard error was calculated from the resampled data.

## Acknowledgements

This work was supported by Welcome-DBT grant and IISER Pune.

## Author contributions

NS: Conceptualization, Methodology, Software, Validation, Formal Analysis, Investigation, Data Curation, Writing original Draft, Visualization, Funding acquisition. TB: Conceptualization, Methodology, Software, Formal Analysis, Investigation, Writing review and editing, Visualization. HL: Conceptualization, Resources, Writing review and editing, Funding acquisition. TS: Conceptualization, Resources, Writing review and editing, Funding acquisition. SN: Conceptualization, Methodology, Software, Validation, Formal Analysis, Resources, Investigation, Writing Original Draft, Visualization, Supervision, Project administration, Funding acquisition

## Competing interests

The authors declare no competing interests.

